# Membrane Transport of Rhodamine 6G by the reconstituted multidrug resistance–ABC transporter Pdr5 in Horizontal Lipid Bilayer

**DOI:** 10.1101/2025.07.31.667618

**Authors:** Daniel Blum, Lea-Marie Nentwig, Stefanie L. Gala Marti, Lutz Schmitt, Richard Wagner

## Abstract

Pdr5 is the most abundant ABC transporter in *Saccharomyces cerevisiae* and plays a key role in the pleiotropic drug resistance (PDR) network, actively preventing the sustained entry of a wide range of structurally diverse compounds into the cell. This transporter has been extensively studied *in vivo*, in plasma membrane vesicles, and more recently, after reconstitution into artificial membranes, making it a widely used model system for ABC transporter research. Recent structural studies of Pdr5 have provided new insights into its architecture and function. However, the precise molecular mechanisms underlying Pdr5-mediated membrane transport remain largely unresolved. In this study, we investigate the interaction of reconstituted Pdr5 in an artificial freestanding horizontal lipid bilayer system with Rhodamine 6G (R6G), a well-known Pdr5 substrate. We show that functionally reconstituted Pdr5 actively transports the R6G^+^ cation across the membrane in an ATP-dependent manner, moving it from one bulk phase to the other.

## Introduction

The ATP-binding cassette (ABC) transporter family comprises one of the important superfamilies of membrane transport proteins and occurs in all areas of life (*1, 2*). With 42,000 copies per cell, Pdr5 is the most abundant ABC transporter in Saccharomyces cerevisiae(*3*) and plays a major role in the pleiotropic drug resistance (PDR) network(*4*), preventing actively sustainable cell entry of many structurally unrelated compounds (*5*). Due to its molecular architecture, Pdr5 serves as a perfect model system for the family of asymmetric ABC transporter (*6*). Important advances in the investigation of the molecular mechanisms of substrate transport by Pdr5 were the functional isolation of Pdr5 and its functional reconstitution in artificial lipid membranes (*7*). Although recently many structures of ABC transporters including Pdr5 have been published (*2*), it remains elusive how these transporters function. In particular, the details on how the energy derived from NTP hydrolysis is converted into mechanical work (e.g.-conformational transitions, upward energetic transport, ATP vs. GTP, co-transport of protons). The fluorescence dye Rhodamine-6G (R6G) has been shown to be a substrate of Pdr5 (*8, 9*). The spectroscopical and ionic properties of R6G making the fluorescent dye an interesting molecular tool for studying the molecular mechanisms of Pdr5 transport for charged substrates (*8, 9*). Classical fluorescence spectroscopy and electrophysiological techniques have already been used to investigate the R6G^+^ transport properties of the multidrug Pdr5 ABC transporter(*9, 10*). Here we describe the investigation of the interaction of the fluorescence dye Rhodamine-6G (R6G) with reconstituted Pdr5 in the artificial horizontal lipid bilayer (HLB)-system using the high-resolution scanning time-correlated single photon counting (TCSPC) technique delivering fluorescence lifetime and fluorescence-intensity data with high spatial 3D resolution (*11, 12*). Using the experimental HLB setup, we studied the interaction between rhodamine 6G (R6G^+^) and the horizontal bilayer as well as bilayers containing the reconstituted multidrug ABC transporter Pdr5 in comparative experiments. In x-z scans across the HLB with 1μm stepping, we determined both the fluorescence lifetime of R6G^+^ at spatially resolved single spots at various z-distances from the bilayer and the fluorescence intensity in the x-z-plane of the HLB obtained by x-z line scans converted into grayscale intensity images as an 2D-resolved indirect measure of the R6G^+^ concentration in the x-z plane within, above and below the free-standing horizontal bilayer (see Figure 1).

**Figure 1:**
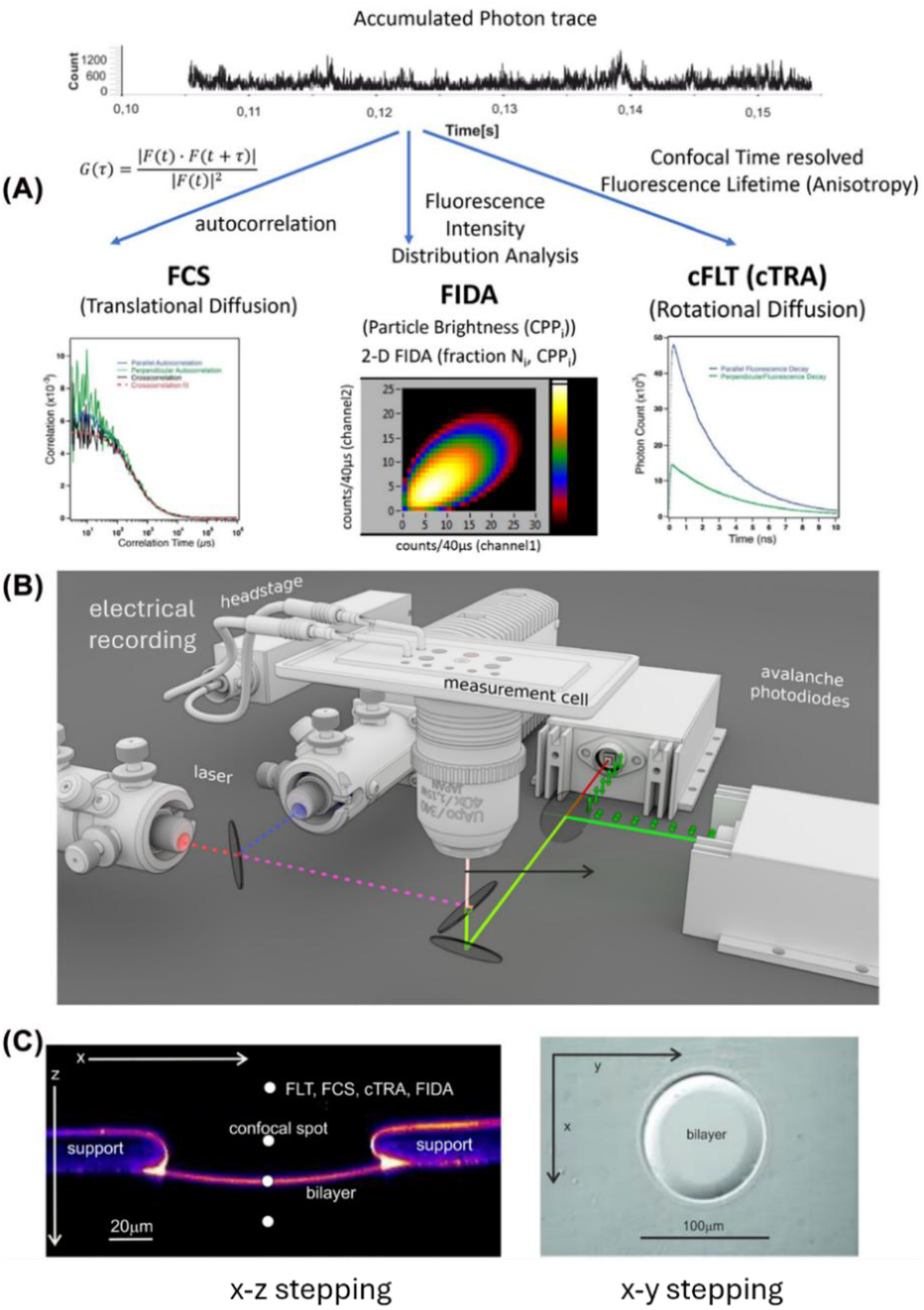
Principal features of the TCSPC technique in the HLB-system. Principal features of the scanning time correlated single photon counting (TCSPC) technique applied to the HLB-system are outlined in Figure 1. TCSPC technique delivers data on (i) fluorescence correlation (FCS) or cross correlation (FCCS), (ii) fluorescence intensity distribution and (iii) fluorescence lifetime as well as fluorescence anisotropy data simultaneously. **(A)** Schematic overview of the principal setup and experimental procedures for the recording of time-correlated-single photon events (top) and their analysis using fluorescence correlation spectroscopy (FCS) (left), analysis of fluorescence intensity distribution (FIDA) (middle) and confocal time-resolved-fluorescence-lifetime anisotropy (cFLT, cTRA) (right). **(B)** Experimental setup of the - confocal-scanning single-photon-counting spectrometer with simultaneous single-channel electrical recording. **(C)** Converted X–Z scan of a fluorescent doted free-standing horizontal bilayer with 50 nm x-z-stepping images (left) and X–Y scan with 200 nm X-Y-stepping images (right).

## Material and Methods

Pdr5 was expressed and purified as described (*7, 13*). Reconstitution of purified Pdr5 into the lipid bilayer was performed as given in detail (*10*).

The horizontal lipid bilayer (HLB) setup as well as details on the electrical and fluorescence recordings from HLB are described in principle and detail elsewhere (*11, 14-16*). In brief, confocal imaging and fluorescence fluctuation recordings were performed on an Insight Cell 3D microscope from Evotec technologies (Hamburg, Germany, now Perkin Elmer), equipped with a 488-nm pulsed diode laser (∼80-ps pulse width; PicoQuant, Berlin, Germany). The signal from each of the two detector units was split up on the correlator and the imaging unit of the Insight and on a PHR 800 router in combination with a PicoHarp 300 counting module (PicoQuant). The PicoHarp 300 allowed interactive analysis, for example the recording of raw photon traces and online FCS and lifetime analysis. The repetition rate of the laser was set to 40 MHz and the resolution of the PicoHarp 300 to 16 ps. For z-scan spot-measurements, the focus was moved in the Z-direction in either 100-nm or 1μm steps through the membrane plane and 20-s photon traces were recorded at each position. The fluorescence intensity within the lipid bilayer plane as well as the cis and trans compartment were visualized with the <u>M</u>odular-Imaging and <u>P</u>hoton-<u>S</u>tatistics-<u>S</u>ystem (MIPSS, Evotec) using stacks of x-z-scans, whereby the spatially resolved 2D-fluorescence intensity in the HLB-system is converted into a gray-value-intensity image. The gray value intensity images were analyzed using the NIH-imaging-software ImageJ (*17, 18*)

### Analysis of grayscale intensity-images obtained by x-z line scans of the fluorescence intensity

Stacks of x-z-scans with the spatially resolved 2D-fluorescence intensity of the HLB-system were converted into a gray-value-intensity image using the <u>M</u>odular-Imaging and <u>P</u>hoton-<u>S</u>tatistics-<u>S</u>ystem (MIPSS, Evotec).

The grayscale value (or intensity) of a fluorescence image is correlated with the concentration of the fluorescent dye in a nonlinear way due to several factors, however at low concentrations, the fluorescence intensity (gray-value) is proportional to the concentration of the dye:

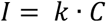

where: I=fluorescence intensity (gray value), C=concentration of the fluorescent dye, k = a proportionality constant that depends on factors like quantum yield, excitation efficiency, and detail of the instrumental setup. It has been shown previously that in the used concentration range of R6G^+^ deviations from linearity that occur due to nonlinear effects at higher concentrations (e.g. inner filter effect, self-quenching) are negligible (*19*). Therefore, the chosen method for analyzing the spatial distribution of the R6G^+^ concentration in the HLB-system can be a useful method with high spatial resolution that allows new important insights into the mechanism of R6G^+^ transport by Pdr5 since it allows an indirect measure of the R6G^+^ concentration in the x-z plane within, above and below the free-standing horizontal bilayer (see Figure 1C).

The grayscale intensity-images obtained from confocal fluorescence intensity x-z-line scans (z-stepping 1μm) were analyzed applying the NIH imaging software ImageJ (*17, 18*). For determination of the R6G^+^concentration in the cis and trans compartment we used the integrated density of gray-values from defined areas of identical size above and below the bilayer. Distances in the 2-D images were calibrated using the known diameter of the *Teflon-Septum-Hole* (typically, d=150μm). The resolution of the bilayer plane was limited by the individual pixel size determined by the number of pixel/μm which was typically between N≅0.5 (pixel/μm) to N≅2 (pixel/μm) corresponding to *F*_*pixel*_ ≅ (1.5*μm*) ^2^to *F*_*pixel*_ ≅ (0.48*μm*)^2^. The area of the bilayer was therefore approximated by a series of individual pixels over the entire bilayer diameter when determining the integrated density of the gray-value. The integrated density of the gray-value intensities results from the product of the average intensity and the size of the respective area:

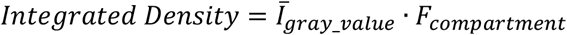

For the identically sized areas above and below the bilayer, typically 25% of the total area of the image were considered.

## Results

We used the TCSPC technique (Figure 1) to perform two main types of measurements on free-standing horizontal bilayers (HLBs): (i) confocal fluorescence intensity x-z-line scans (z-stepping 1μm) and (ii) confocal single point x-z-spot lifetime scans (z-stepping 0.1μm). The gray-scale-intensity images obtained from the x-z line scans were analyzed using *Image J* (*20*). To optimize the visual appearance of the images shown in the following and above in Figure 1c, the grayscale intensity images from the x-z line scans were converted into pseudo color Images using the *Image J Lookup Table “Fire”* (*20*). Individual confocal fluorescence intensity images obtained by x-z line scans were calibrated to the respective laser excitation energy. In the following analysis of our data, we use the calibrated gray-value-intensity of the different images as an indirect, relative measure of the concentration of the fluorophore R6G^+^ along the z-axis and in the x-z planes in the different compartments (cis/bilayer/trans) of the HLB system. In addition, we use the concentration dependence of the spatially resolved fluorescence lifetime (τ_*F*_) as an analysis tool to obtain spatial resolved values of the R6G^+^ concentration in close vicinity of the bilayer in the HLB-system along the z-axis.

### Control Bilayer

In order to improve the visual appearance, the grayscale intensity-images obtained from confocal fluorescence-intensity x-z-line scans (z-stepping 1μm) were converted by the color lookup table *Fire* of the ImageJ software (*17, 18*) that can be applied to grayscale images to produce pseudo color Images.

Figure 2 top shows example images of the x-z converted line scan gray-scale-intensity recording in a control HLB (top left) and recordings from bilayer after addition of 1μM R6G^+^ at t=0 to the cis compartment at the given time increments after R6G^+^ addition. The corresponding gray-scale-intensity values of the intensity profile along the depicted lines in z-direction are shown below.

**Figure 2:**
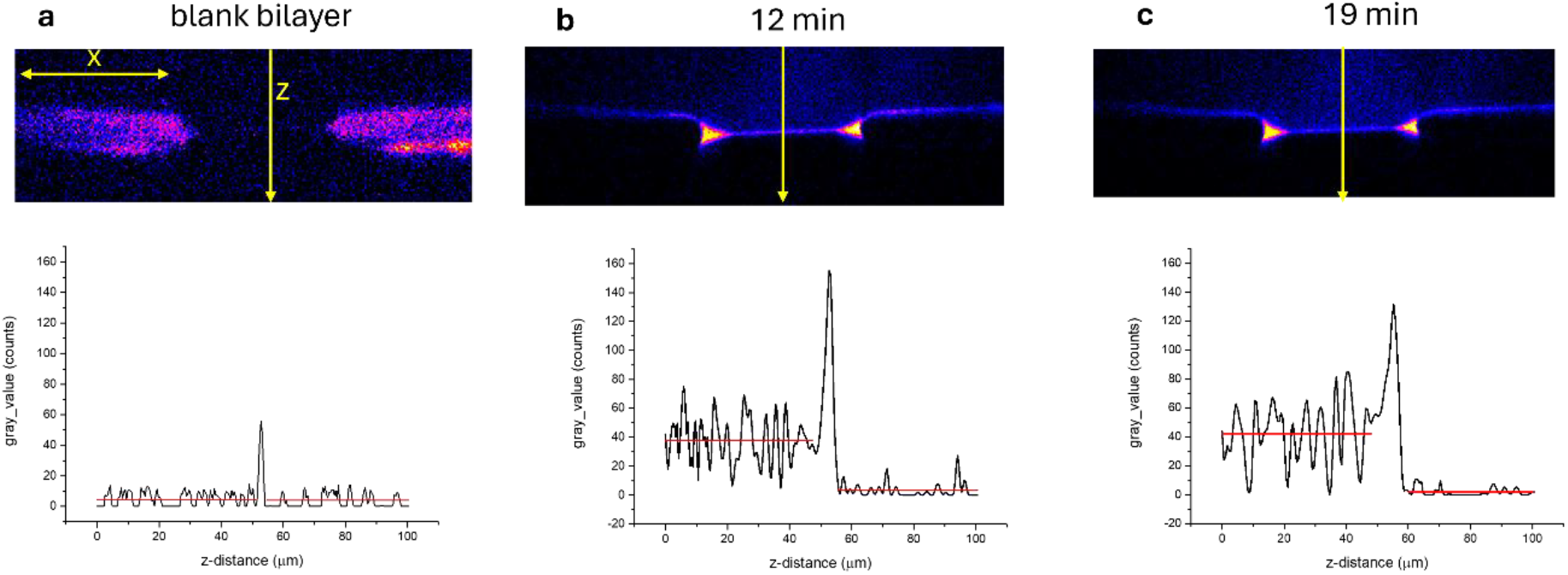
**Top:** x-z scan recording from the control HLB **(a)** and recordings after addition of 1μM R6G^+^ final concentration at t=0 to the cis compartment at 12min **(b)** and 19min **(c)** time increments after R6G^+^ addition. **Lower:** Profile of the gray-value intensity along the z-axis (average of 3-scans). Red lines show the mean gray values.

A more detailed analysis of data as shown in Figure 2 is depicted in Figure 3. Grey-value intensity along the z-axes (z-scan) of the HLB-system is shown as an example in Figure 3a. This results in the mean values for gray-value-intensity for the cis, trans compartments and the bilayer from several measurements (Figure 3b). The integrated gray-value-density representing an indirect measure of the R6G^+^-fluorophore distribution within the 3-compartments of the HLB is shown in Figure 3c. Figure 3d shows a comparison of the area referenced mean-gray-value/μm^2^ of the cis-plane and the bilayer plane.

**Figure 3:**
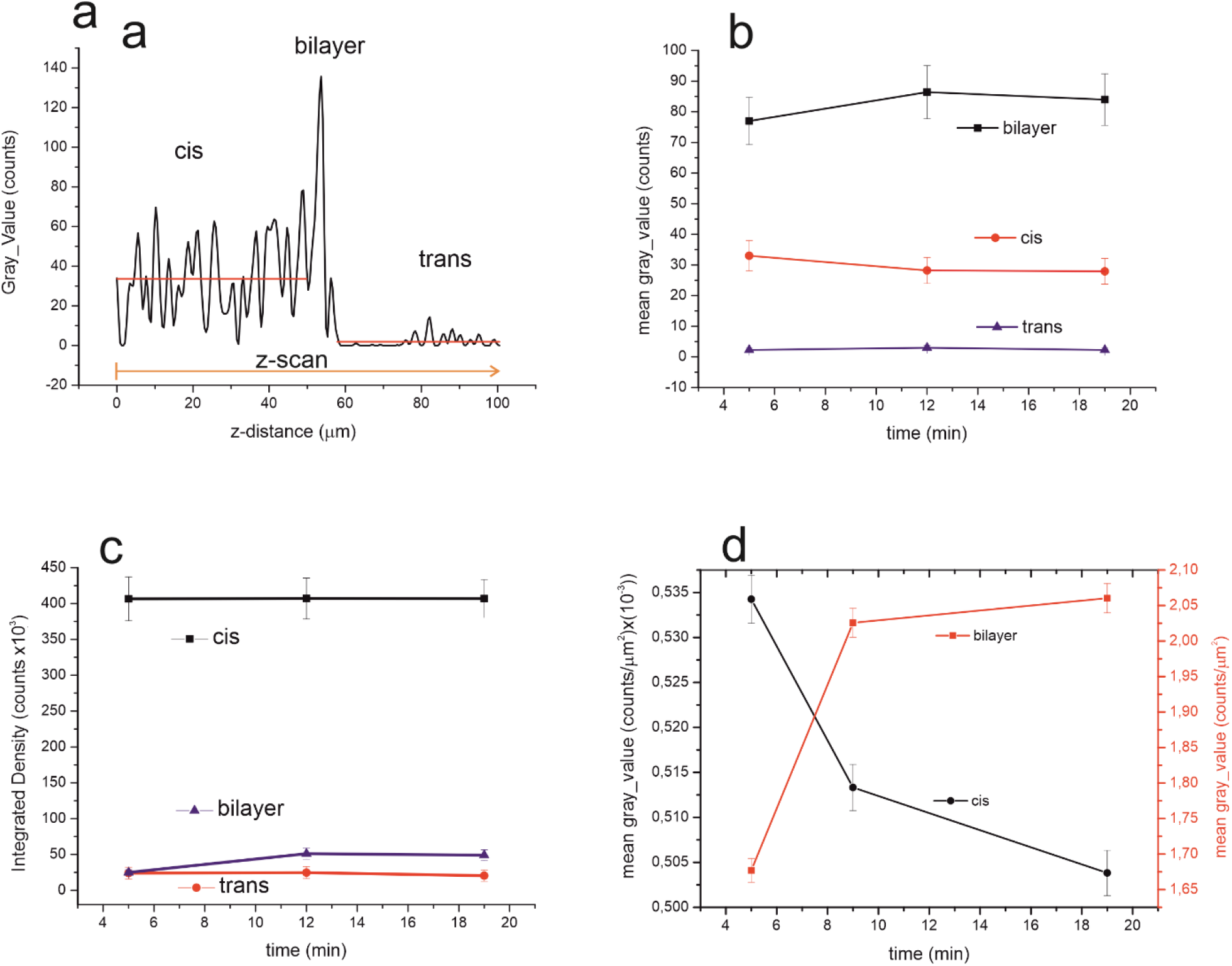
**(a)** Gray value counts along the z-axis of the HLB, average of 3 z-profile scans, straight red lines mean gray values cis and trans. z-scan profiles from the image at t=5min after R6G addition (see also Figure 2b, c). **(b)** Averaged gray values of 5 z-scan-profiles 5min, 12min and 19min after addition of R6G^+^ **(c):** Integrated density of gray value counts in the cis, trans compartment and the bilayer plane of HLB bilayer after addition of cis 1μM R6G^+^ at t=0min. **(d)** Integrated density of gray value counts per μm^2^ in the cis, compartment and the bilayer plane of HLB bilayer after addition of cis 1μM R6G^+^ at t=0min.

The mean gray value data from the z scan profiles at different times after the addition of 1μM R6G^+^ to the cis compartment (Figure 2a, b, Figure 3a, b) demonstrate that the bilayer very effectively separates the cis und trans compartment from each other and the R6G^+^ cation cannot permeate across the membrane. Similarly, the evaluation of integrated gray-value-density of the x-z scans (Figures 3d) above and below the bilayer reveals that the membrane is not permeable to the R6G^+^ cation even after a prolonged period the R6G^+^ concentration in the cis compartment remains almost constant while in the trans compartment the integrated density values of the trans compartment remained constant at the blank level. However, the integrated density levels in the bilayer plane increased, indicating that R6G^+^ enriches significantly in the bilayer. Comparing the integrated density of the gray-value-intensity of the cis compartment to the integrated gray-value-density of the bilayer plane, enrichment of R6G^+^ in the bilayer-plane reaches a factor of f_enr_≅8.2 (Figure 3c) although R6G^+^ in the present buffer solution of the cis compartment (250 mM KCl, 10 mM HEPES, pH 7.0) is predominantly present as a cation R6G^+^ (*21*). However and most important, in terms of concentration changes, corresponding in the 2-D case to the average grey-value-intensity per μm^2^, -this means that the concentration of R6G^+^ in the bilayer membrane is more than 3 orders of magnitude higher (Δ*c*_*bilayer*_ ≅ 4100 μ*M*) than in the start concentration in the cis-compartment 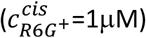 (Figure 3d). This concentration-ratio would correspond to a partition coefficient of 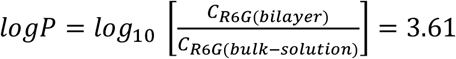.

To gain further basic information on the fluorescence-spectroscopic properties of R6G^+^ in aqueous buffer solution (250 mM KCl, 10 mM HEPES, pH 7.0) and particularly at the bilayer membrane surface we first determined as a reference the fluorescence lifetime of R6G^+^ in the same isotropic buffer solution as applied in the HLB-experiments over a wide concentration range (0.01M to 10^−8^M) (Figure 4a,b).

**Figure 4:**
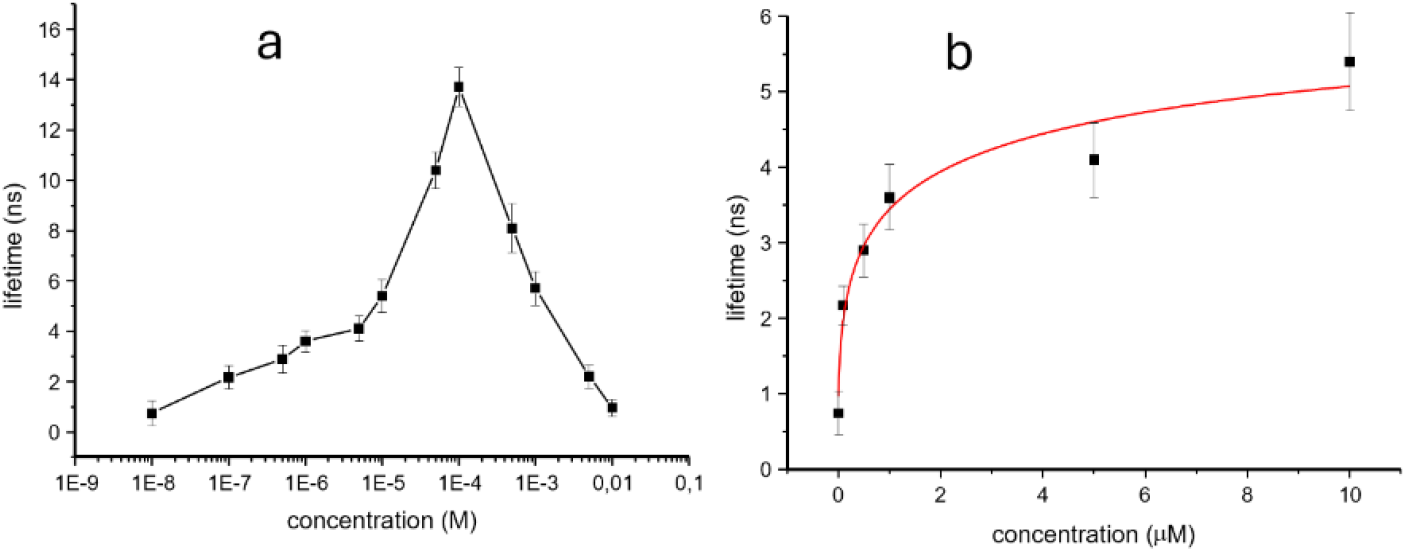
Concentration dependence of the R6G fluorescence lifetime in aqueous buffer solution (250 mM KCl, 10 mM Hepes, pH 7.0) **(a)** concentration range log-scale 0.01M to 10^−8^M. **(b)** concentration range 0.01μM to 10μM

As observed previously for different organic solvents (*22, 23*) the lifetime of R6G in the aqueous buffer solution is significantly dependent on the concentration. Increasing from τ_*F*_ *≅*1 *ns* (10nM) up to τ_*F*_ *≅*14 *ns* (100μM) while dropping to τ_*F*_ *≅*0.7 *ns* again at a concentration of 10mM (Figure 4a). In the concentration range between 1μM and 5μM R6G^+^, which is relevant for the experiments described below, τ_*F*_ increases exponentially from τ_*F*_ *≅*1ns (1μM) to an asymptotic value of τ_*F*_ *≅*5*ns* (10μM) (Figure 4b).

The concentrations of R6G^+^ used during the experiments in the aqueous buffer solution of the cis-compartment the HLB-system (1μM to 5μM) was well below the concentration at which higher aggregates i.e. trimerization occurs so that in this case only monomer–dimer equilibrium needs to be considered (*24*). The value of dimerization constant 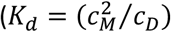 of R6G^+^ was determined to be K_d_=5.75× 10^3^ (*21*), thus in our applied concentration range of R6G^+^ in the HLB-experiments the concentration of the dimer is negligible low (*21, 24*) and need not be taken into account in the further calculations and discussions.

In the further course we determined the fluorescence lifetime in the HLB system in defined z-spot scans (fixed x-coordinate) and z-increments with 1μm z-stepping (z=±10μm) above the bilayer-plane (cis), in the bilayer plane and below the bilayer plane (trans) using the spatially resolved TCSPC-technique. Measurements were performed after adding a final concentration of 1 μM R6G^+^ to the cis compartment starting at t_addition_≥10min at a fixed x-coordinate in z=1μm steps. The spatial resolution of the z-spot scanning lifetime measurements is mainly limited by the geometry of the confocal volume (*25-27*) in particular the axial (z) height of the confocal volume which has been determined for the used setup to be lateral (xy) width d_xy_ ≅600nm and axial (z) height h_z_ ≅1.2μ) (*28*) thus limiting the resolution of z-point scans.

Fluorescence lifetime data as shown in representative Figure 5 were obtained from reference control bilayer and bilayer containing the reconstituted Pdr5 with additionally 5mM ATP, 1mM Mg^2+^ in the cis compartment.

**Figure 5:**
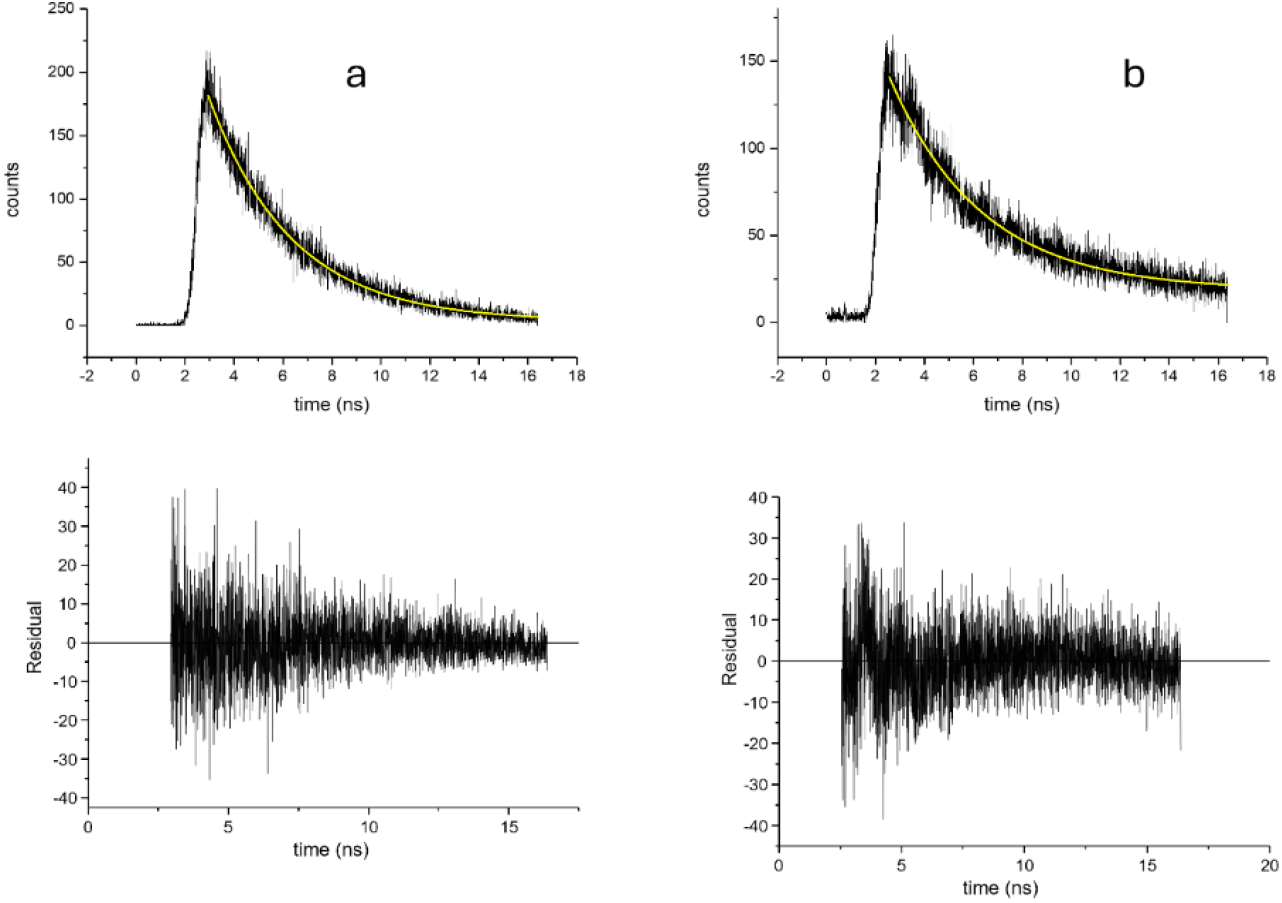
Examples of fluorescence-decay curves from a control bilayer (x=20μm, z=0μm) **(a)** and **(b)** a bilayer containing the reconstituted Pdr5 and additionally 5mM ATP, 1mM Mg^2+^ (x=20μm, z=-1μm).

The values obtained from Figure 5 were τ_*F*_ =3.4ns for **(a)** (control panel) and τ_*F*_ =3.8ns for **(b)** (Pdr5-containing bilayer). The summary of the fluorescence lifetime measurements with control bilayers are shown in Figure 6a, b.

**Figure 6:**
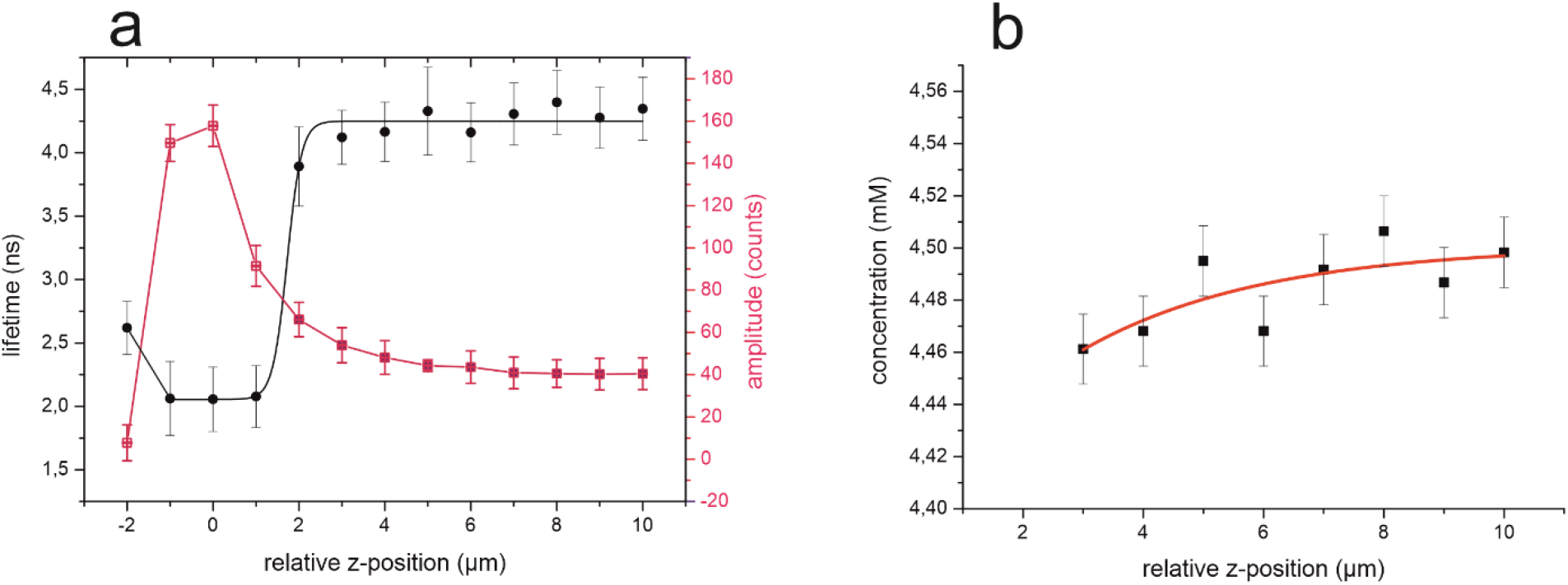
**(a)** Lifetimes and the corresponding fluorescence intensity amplitudes obtained from decay curves as shown in Figure 5 along the z-coordinate relative to the location of the bilayer (z=0μm). **(b)** Lifetime converted into the concentration along the z-axis (data from n≥3 measurements) for details see text.

Figure 6a shows the lifetime and the corresponding fluorescence amplitude obtained from the decay curves along the z-coordinate relative to the location of the bilayer (z=0μm). The z-coordinate of the confocal volume was moved at a fixed x-coordinate in 1μm steps above the bilayer (+z) and below the bilayer (−z) (see also Figure 1). It is important to note that below the bilayer in the trans compartment at z≤-2μm no fluorescence decay signal could be observed furthermore. As obvious from Figure 6a the lifetime from z=-2μm decreased first from τ_*F*_ = 2.6*ns* to τ_*F*_ = 2.0*ns* in the bilayer plane (z=-1 to z=+1) followed by an asymptotic increase to τ_*F*_ *≅*4.3*ns* at z=+10μm (cis compartment) (Figure 6a). In isotropic buffer (250 mM KCl, 10 mM HEPES, pH 7.0) solution of 1μM R6G^+^ the observed lifetime was τ_*F*_ = 3.34 ± 0.03*ns*. The amplitude plot in Figure 6a where the amplitude within the bilayer plane is about 4fold increased compared to the cis solution also supports a significantly higher concentration of the R6G^+^-dye within the bilayer plane (−2μm ≤z ≥+2μm) as compared to the cis solution.

Assuming that the fluorescence properties of the R6G^+^-dye in isotropic solution and those above the bilayer-plane (z=+3 to z=+10) the cis compartment are in a reasonable approximation comparable, the fluorescence lifetime along the z-axis of the HLB system close to the bilayer-plane (z≥+1μm) in the cis compartment can be converted into the concentration profile along the z-axis using the calibration of R6G^+^-lifetime versus concentration (Figure 4b). Using this calibration curve from Figure 4b we converted the lifetime into concentration values and plotted the data against the z-coordinate of the HLB (Figure 6b). This plot indicates that the concentration of R6G in the cis compartment directly above the bilayer (z=+3 to z=+10) is about 4-fold higher than the applied final R6G^+^ concentration of 1μM to the cis compartment. From this it can be concluded that after adding R6G^+^ to the cis compartment, a concentration gradient formed in the HLB system in which the local concentration of R6G directly above the bilayer (z=+3 to z=+10) is approximately 4 times higher than in the bulk solution (z≤+10).

Within the z-coordinates -1μm ≤z ≥+1μm, the bilayer plane, τ_*F*_ decreased significantly. Considering that the local R6G^+^ concentration in the bilayer-plane is presumably about 3 orders of magnitude higher than in the bulk solution (Figure 3d), the fluorescence lifetime values also show that the R6G^+^concentration in the bilayer-plane reaches values in the region of 1mM. This value is in the same order of magnitude as gained from the analysis of the gray-scale-intensity-images obtained from confocal fluorescence-intensity x-z-line scans (see Figure 3d). We will discuss this topic in more detail below.

Here we want to highlight that in the HLB the fluorescence intensity at the coordinate z≤-2 dropped below the threshold value where no fluorescence signal could be observed anymore (≤10^−14^M R6G^+^, (*29*)) showing that no detectable amounts of the dye were present in the trans-compartment.

The fact that the bilayer in these lifetime spot-z-scans does not form a similar sharp boundary as observed in the images generated by x-z line scans (e.g. Figure 2) is certainly due to the geometry of the confocal volume (*25*). The longer z-axis of the prolate shaped ellipsoidal confocal volume of the experimental set up was determined to be *h*_*z*_*≅*1.2*μm* (*30*). Thus, the spatial resolution of the z-point-scans are limited by this factor. While within x-z-line scans overlapping of z-values occurs resulting in slightly increased resolution (*26, 27, 31*).

### Pdr5 Bilayer

To monitor the incorporation of Pdr5 into the HLB during reconstitution we measured routinely the electrical conductance of the bilayer before and after Pdr5 reconstitution. A representative example is shown in Figure 7.

**Figure 7:**
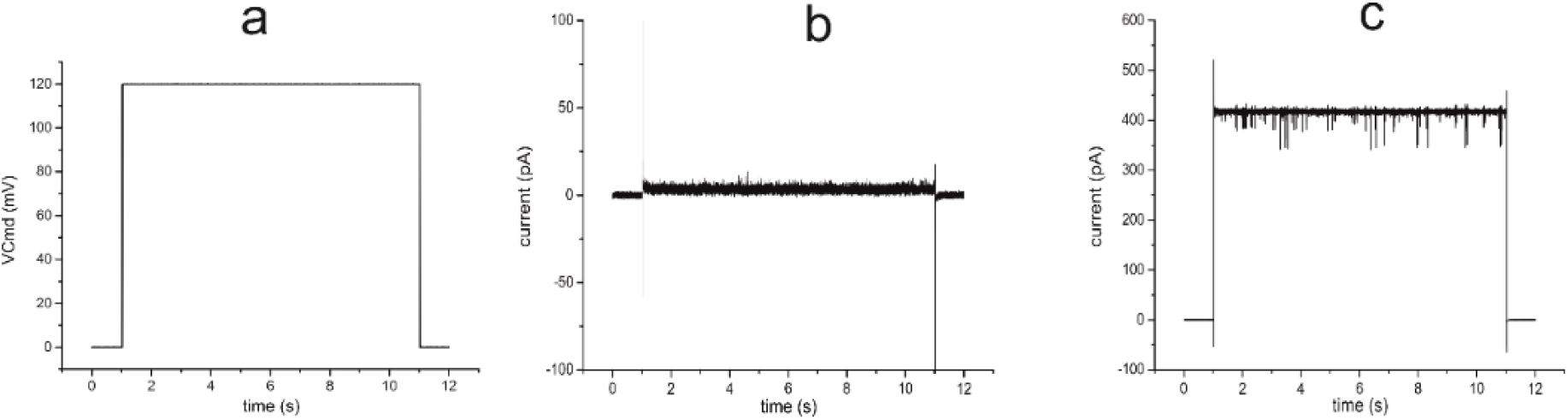
**(a)** Voltage-gate applied to the HLB. Electrical recording from the HLB before **(b)** and after reconstitution of Pdr5 **(c)**.

Figure 7a shows the applied voltage gate (V_cmd_=+120mV) during the voltage clamp recording. The control bilayer revealed a low basic conductance of 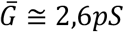 (Figure 7a) a value typically observed for non-conducting lipid bilayer, while after Pdr5 reconstitution the observed average conductance was 3 orders of magnitude higher with 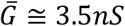 (Figure 7c), revealing only brief channel closing gating events with conductance values of 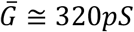 and 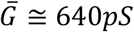 respectively. The closure gating events are in the same order of magnitude as observed previously for the activated Pdr5 channel (*32*). There is good evidence that the majority of the current in Figure 7c is flowing through a significant, larger number of reconstituted non-activated Pdr5 channel pores (*32*). Using the deduced value of the non-activated Pdr5 channel of 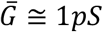 (supplement in reference (*32*)) this would mean that about 3500 non-activated Pdr5 were involved in the observed ion conduction (Figure7c). In summary the electrical recordings from the HLB showed that Pdr5 was successfully reconstituted into the HLB.

Next, we analyzed representative gray-scale-intensity images of the HLB-system obtained by x-z line scans after addition of 1μM R6G^+^ cis to a bilayer containing the reconstituted Pdr5 in the presence of 2mM ATP, 2mM Mg^2+^ in the cis compartment (Figure 8). The x-z line scan recordings were performed at the given time intervals after addition of 1mM R6G^+^ to the cis compartment (see Legend).

**Figure 8:**
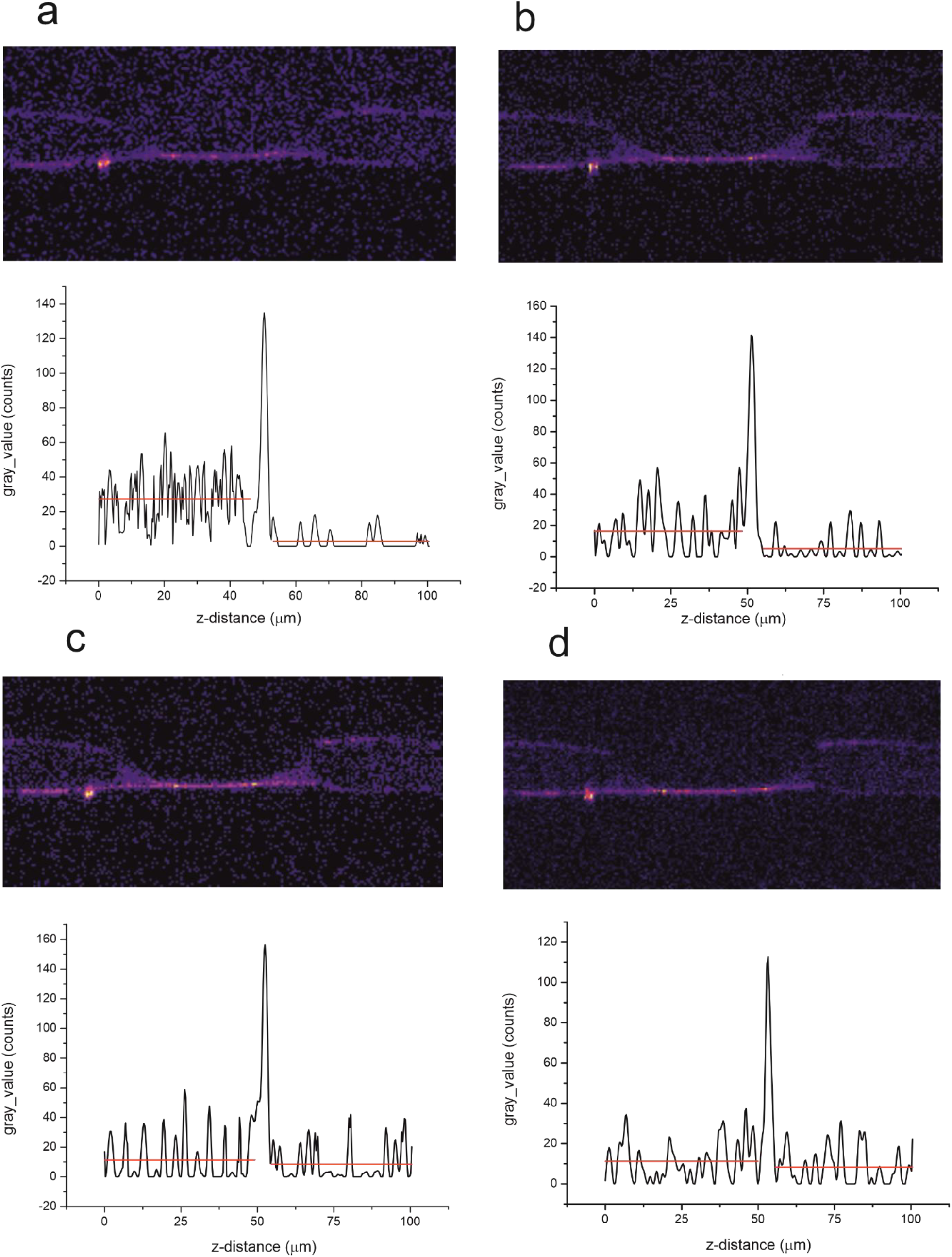
Representative converted gray-scale intensity images of the HLB-system obtained by x-z line scans after addition (t=0min) of 1μM R6G^+^ to the cis compartment of the bilayer containing the reconstituted Pdr5 in the presence of 2mM ATP, 2mM Mg^2+^ in the cis compartment. x-z line scan recordings were performed at t=4min (a), t=5min (b), t=9min (c), t=19min (d) -after R6G^+^ addition. Below the images the corresponding z-profiles of the gray-scale-intensity is shown (average of 3 profiles each).

The Images obtained by x-z line scans were calibrated to the respective laser excitation energy and the obtained gray-scale-intensity images were analyzed with respect to the integrated density gray-scale-intensity values of the 3 different HLB-compartment x-z areas (cis/bilayer/trans). The area of the bilayer was approximated by a series of individual pixels (pixels size typically *F*_*pixel*_ ≅ (0.5*μm*)^2^(see Methods for details). The integrated density of the gray-value intensities is the product of the average gray-value-intensity and the size of the respective areas.

The time dependent changes of the integrated density of gray-value-intensities describe qualitatively the temporal changes in the distribution of R6G^+^ between the three compartments of the HLB system. While in the control bilayer (Figure 3c) it was only observed that R6G^+^ accumulates largely in the membrane, the integrated density of gray-value-intensities in the cis compartment changes only marginally and in the trans compartment the integrated density of gray-value-intensities values remain constantly low at the background level. Analogous results were observed when the cis compartment contained no ATP/Mg^2+^. However, in the presence of Pdr5 in the membrane and ATP/Mg^2+^ cis, the distribution of R6G^+^ between the 3 HLB-compartments changes drastically. The values for the integrated density of gray-value-intensities in cis decrease exponentially and the values in trans increase accordingly exponentially (Figure 9a). The same is observed for the bilayer-plane where the values for the integrated density of gray-value-intensities also increase exponentially (Figure 9b). For the values relating to the bilayer, however, it is important to consider that the increase in the R6G^+^ concentration in the membrane after the addition of 1μM R6G^+^ to the cis compartment the incorporation is initially so rapid (t≤ 1min) that this part of the uptake process could not be resolved experimentally in terms of time.

**Figure 9:**
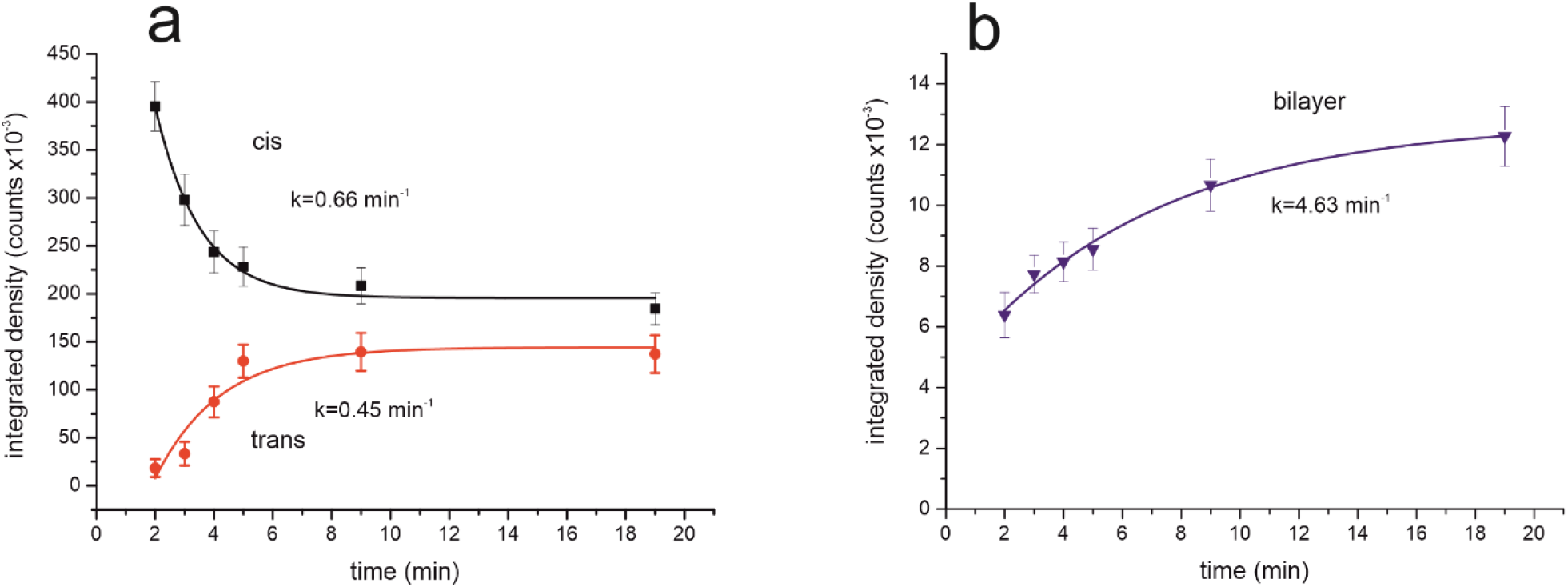
Time dependence of the integrated density of gray-value-intensities cis/trans **(a)** and the bilayer **(b)** after addition of 1μm R6G^+^ at t=0min to the cis compartment (averages from 2 x-z-line scan gray-value-intensity images).

In summary the above results show that after reconstitution of Pdr5 into the horizontal bilayer in the presence of 1μM R6G^+^, 5mM ATP, 1mM Mg2^+^ in the cis compartment, -the R6G^+^cation is actively transported across the membrane into the trans compartment. In contrast to this, the control bilayer membrane was completely impermeable to the R6G^+^ cation (Figure 2).

It is worth noting that only in the presence of ATP and Mg^2+^ in the cis compartment a decrease of the R6G^+^intensity in the cis compartment and an increase of the R6G^+^intensity in the trans compartment could be observed.

In a further analysis, we calculated from the x-z line scan-generated gray-value-intensity images the time dependence of the mean gray-values related to the areas (μm^2^) above and below the bilayer as well as the area of the bilayer. In the present 2D case, these values are representative of the time course of the concentration changes of R6G^+^ in the three HLB-compartments after addition of 1μM R6G^+^ at time t=0min. The results of this analysis are shown in Figure 10 a, b.

**Figure 10:**
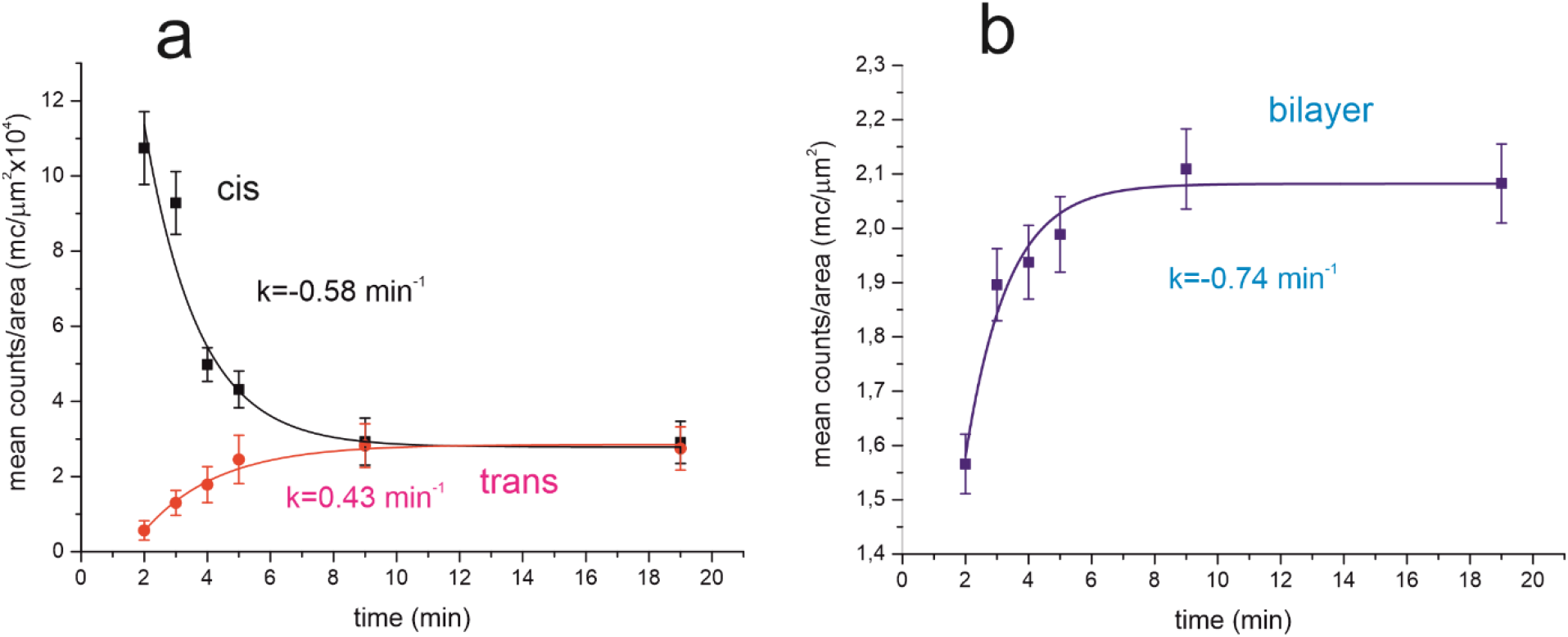
Time course of the mean-gray-value-intensity/μm^2^ of the 3 HLB-compartments (a) cis and trans compartment (b) bilayer after addition of 1μM R6G^+^ at t=0min to the cis compartment (data from n=2 bilayer).

Similarly, as observed for the integrated density of gray-value-intensities analysis above the mean gray value area density of cis compartment decreased exponentially, while the gray value area density of trans compartment increased exponentially. The determined rate constants were rather close to the one observed with the z-Profile analysis above (Table 1). Remarkably, the gray-value area-density of the bilayer shows a significant exponential enrichment of the R6G^+^ dye in the membrane (Figure 10b). For the values relating to the bilayer, however, it is again important to consider that the increase in the R6G^+^ concentration in the membrane containing the reconstituted Pdr5 after the addition of 1μM R6G^+^ to the cis compartment the R6G^+^ incorporation is initially so rapid (t≤ 1min) that this process could not be resolved experimentally in terms of time. Thus, the time course in Figure 10b represents probably only a second slower phase of the process. This also holds for the time dependent decrease of R6G^+^-concentration the cis compartment (Figure10a). While the kinetics of the R6G^+^-concentration increase in the trans compartment is a product of two coupled kinetic processes, the enrichment of R6G^+^ in the membrane and the membrane transport of the dye by the Pdr5 ABC-transporter.

**Table 1.**
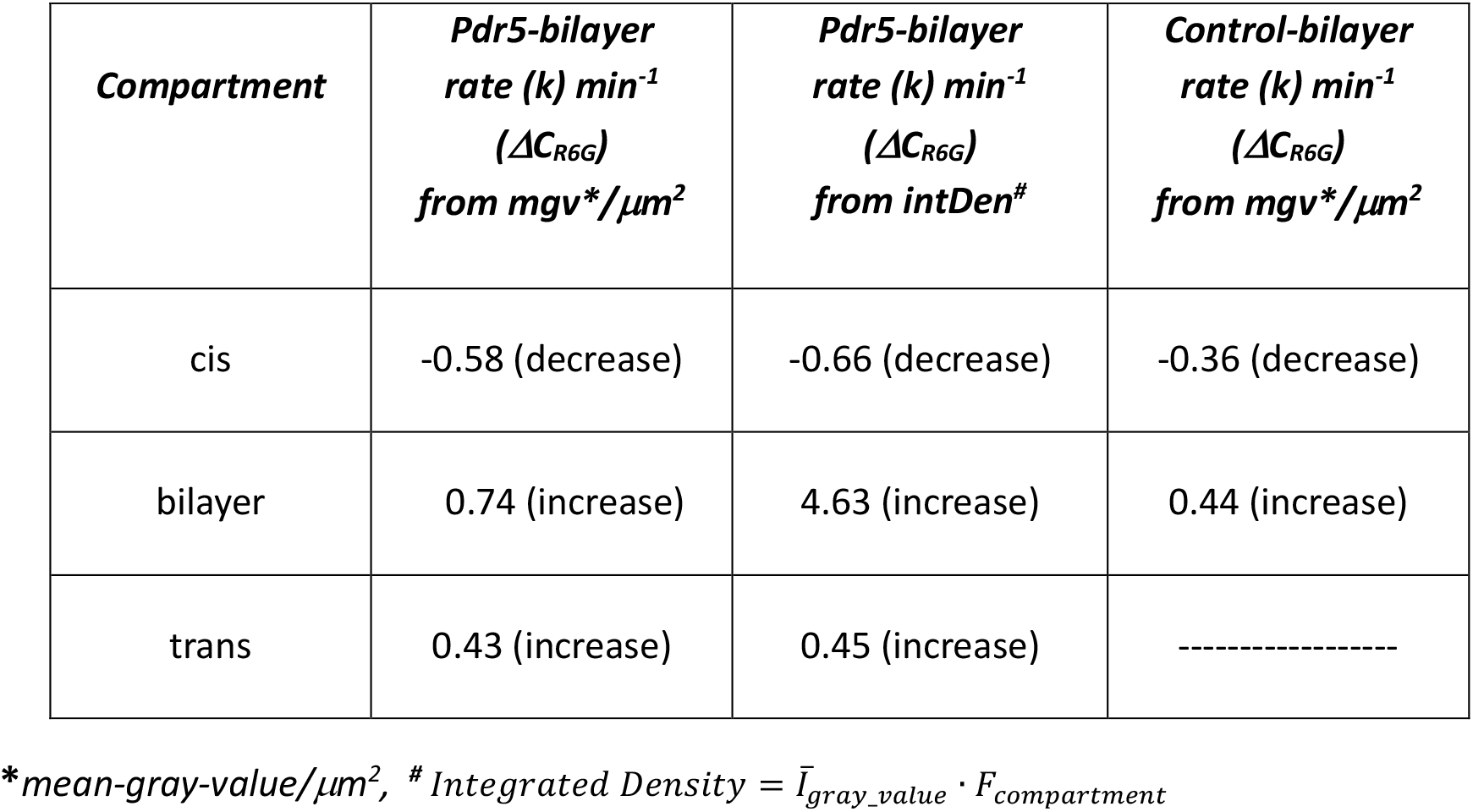
Rate (k) of the R6G^+^-concentration change in the indicated compartment of the HLB for the control bilayer and bilayer containing the reconstituted Pdr5 in the bilayer and 5mM ATP, 1mM Mg^2+^ on the cis side after addition of 1μM R6G^+^ at t=0min to the cis compartment (data from n=2 bilayer).

| **Compartment** | ***Pdr5-bilayer***<br>**rate (k) min<sup>-1</sup>**<br>**(ΔC<sub>R6G</sub>)**<br>**from mgv\*/μm<sup>2</sup>** | ***Pdr5-bilayer***<br>**rate (k) min<sup>-1</sup>**<br>**(ΔC<sub>R6G</sub>)**<br>**from intDen<sup>#</sup>** | ***Control-bilayer***<br>**rate (k) min<sup>-1</sup>**<br>**(ΔC<sub>R6G</sub>)**<br>**from mgv\*/μm<sup>2</sup>** |
| --- | --- | --- | --- |
| cis | -0.58 (decrease) | -0.66 (decrease) | -0.36 (decrease) |
| bilayer | 0.74 (increase) | 4.63 (increase) | 0.44 (increase) |
| trans | 0.43 (increase) | 0.45 (increase) | ----- |

\*mean-gray-value/μm<sup>2</sup>, # Integrated Density = $\bar{I}_{gray\_value} \cdot F_{compartment}$

Remarkable with regard to the kinetic course of the concentration changes of R6G^+^ in the three HLB compartments, visible as changes in the average grey-value-intensity per μm^2^, reveals that the R6G^+^-concentration in the bilayer membrane containing the reconstituted Pdr5 is more than 3 orders of magnitude higher 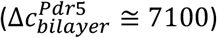 than at the initial start (t=0 min) in the cis-compartment 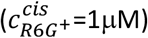 (Figure 10a,b).

A summary of representative fluorescence lifetime measurements from bilayer with 1μM added R6G^+^ cis containing the reconstituted Pdr5 and 5mM ATP, 1mM Mg^2+^ on the cis side are shown in Figure 11.

**Figure 11:**
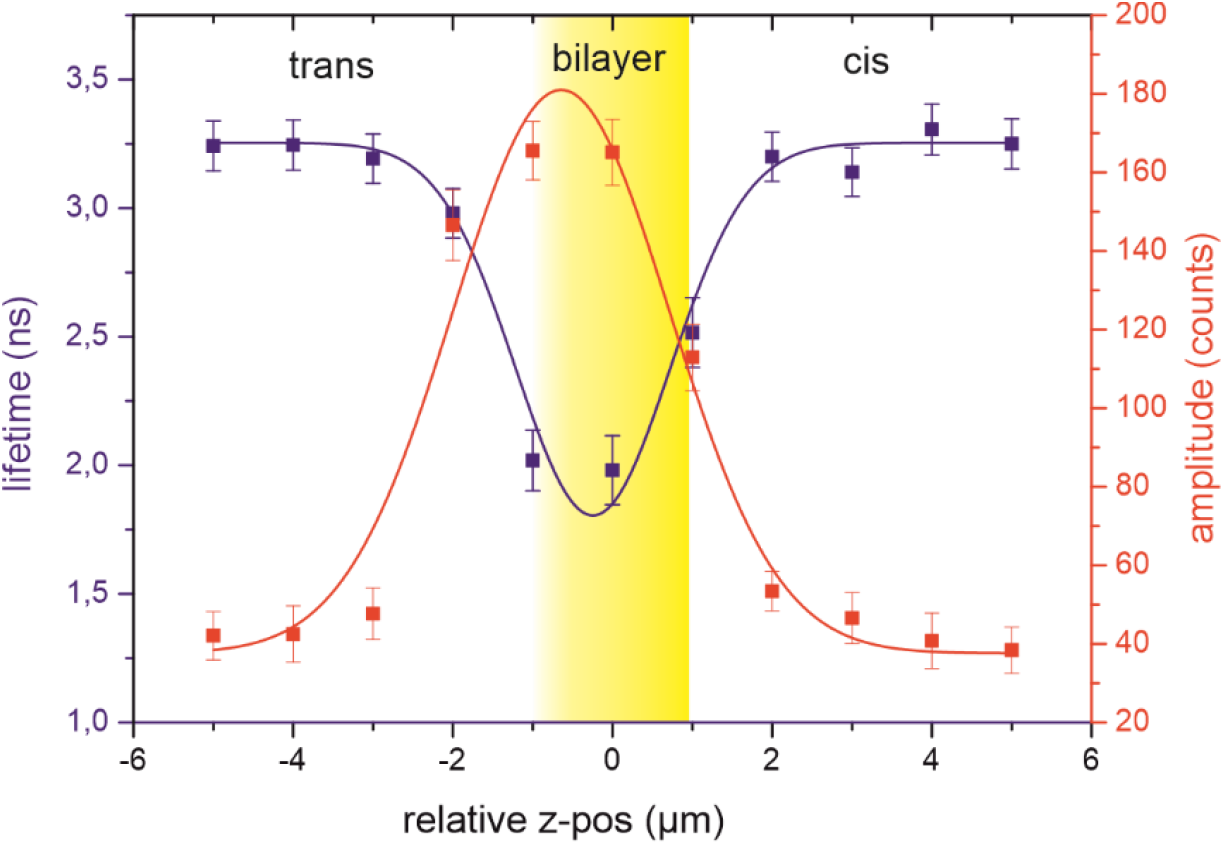
Lifetime and the corresponding fluorescence amplitude obtained from the decay curves along the z-coordinate relative to the location of the Pdr5 containing bilayer (z=0μm) with 1μM added R6G^+^ cis and 5mM ATP, 1mM Mg^2+^ on the cis side.

Figure 11 shows the lifetime and the corresponding fluorescence amplitude of the decay curves along the z-coordinate relative to the location of the bilayer. As obvious from Figure 11 the lifetime of the R6G^+^ fluorescence dye remained on the cis side nearly constant up to the z-coordinate z=+2μm decreasing than slightly to a minimum at z=0μm (the bilayer plane) while increasing again at z=-2μm (trans side) to a similar value as before on the cis side. The amplitudes of the decay-curve fit along the z-coordinate revealed a similar shape as the fit of the lifetime values along the z-coordinate, was however slightly shifted by roughly 1μm towards the trans side of the bilayer. Remarkably, with the reconstituted Pdr5 in the membrane at the HLB z-coordinate z=-3 the amplitude of the fluorescence decay-curves did not drop below threshold values as observed for the control bilayer indicating that significant amounts of the dye were present in the trans-compartment. Again the observation that the bilayer in these point lifetime z-scans does not form a similar sharp boundary as observed in the images generated by x-z line scans in Figures 2 and 7 can be attributed to the geometry of the confocal volume (*25*) with a length of the z-axis of the confocal volume of z ≅1.2μm limiting the resolution of z-point scans.

In this context, it is important to note that the accumulation of R6G^+^ in the membrane plane -deducible from the decreasing lifetime values in the bilayer-plane, -similar to the control bilayer, was presumably also shifted towards the mM regime (Figure 4a) in line with the data of the mean-gray-value-intensity/μm^2^ determinations (Figure 10 a, b).

To obtain further information on the nature of the interaction between R6G^+^ and Pdr5 bilayer, we analyzed the fluorescence anisotropy of R6G^+^ -in solution -at the membrane surface and -in the membrane. The steady state anisotropy (r_0_) and the decay of the anisotropy (r(t)) provide direct information about the mobility and rotational motion of the chromophore in its direct environment (*33-35*). Whereby the possible range of anisotropy values are between r_0_=0.4 (fixed, immobile) and r_0_=0 (high mobility, τ_*rot*_ « 1*ns*) (*34*).

Confocal fluorescence steady state anisotropy measurements in the HLB-system were performed 5min after the addition of 5 µM R6G^+^ to the cis compartment (Table 2) and the steady-state anisotropy (r_(0)_) was determined:

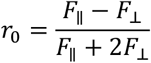

**Table 2.** R6G-Steady state fluorescence anisotropy in different compartments along the HLB-z-coordinate (data from N=3 measurements).

| **Compartment** | **Steady State Anisotropy ($r_0$)** | **Remarks** |
| --- | --- | --- |
| 5 μM R6G<sup>+</sup> in buffer | 0.02 | very high mobility |
| R6G<sup>+</sup> in control Bilayer (z=0μm) | 0.15 | partially restricted mobility |
| R6G<sup>+</sup> in Pdr5 Bilayer (z=0μm) | 0.28 | severely impaired mobility |

These results show that a gradient exists for the rotational mobility of R6G^+^ along the HLB along the z-axis: high mobility in the isotropic solution above the bilayer, restricted mobility at the interface to the bilayer and a severely impaired mobility in the bilayer.

### Summary

We investigated the interaction between the fluorescent dye R6G^+^ with artificial planar bilayer membranes and bilayer containing the multidrug ABC transporter Pdr5 in a horizontal free standing lipid bilayer (HBL) system using the TCSPC technique (*11*). Confocal fluorescence intensity 2D x-z-line scans were used to obtain gray scale images with spatial 2D-resolution of the HLB-cis and trans compartments separated by the artificial planar bilayer. In addition, confocal single point z-spot lifetime scans were performed to obtain spatially resolved information on the fluorescence lifetime along the z-coordinate of the x-z area. Spatial resolved comparative analysis of the image gray-value-intensity of control bilayer and bilayer containing the reconstituted Pdr5 allowed an indirect measure of the concentration of the fluorescence dye R6G^+^ within the three respective HLB-system compartments (cis → bilayer → trans). The data with control-bilayer on the partition of the R6G^+^ between the lipid bilayer and the bulk solution revealed surprisingly a weak hydrophobicity of the cation with *logP* = 3.61. Moreover, after reconstitution of Pdr5 in planar bilayers and addition of 1μM R6G^+^, 5mM ATP, 1mM Mg^2+^ to the cis compartment, we were able to determine the temporal change in the distribution of R6G^+^ between the compartments and their respective rate constants. In addition, 2D-resolved fluorescence lifetimes of confocal spots along the z-coordinate within the HLB-system allowed further spatially resolved determination of the R6G^+^ actual concentration in the three HLB compartments. The study on the interaction between the fluorescent dye R6G^+^ and the multidrug ABC transporter Pdr5 in the horizontal lipid bilayer system revealed significant findings:

- The artificial horizontal bilayer efficiently separates over a longer period of time (≥30 min) the cis from the trans compartment. When R6G^+^ added to the cis compartment in the HLB system, the dye was not permeable through the membrane (Figure 2,3). The analysis of the images generated by confocal x-z line-scans showed that the fluorophore R6G^+^ accumulates more than three orders of magnitude (≅4100times) at the membrane surface plane as compared to the initial bulk concentration (Figure 2, 3d).
- Measurements of the fluorescence lifetime of R6G^+^ along the z-coordinate in the HLB control bilayer show that after addition of 1μM R6G^+^ cis -a concentration gradient is formed such that directly above the bilayer (z=+3μm to z=+10μm) the R6G^+^ concentration was about 4.5 times higher as compared to the added final concentration (1μM) (Figure 6b). The decrease of the R6G^+^ fluorescence lifetime within the bilayer-plane (z=-1μm-z=0 μm-z=1μm) indicates also a high R6G^+^ concentration in the mM regime (Figure 6a, Figure 4a) within the bilayer.
- A remarkable result of the investigations in the HLB system was moreover that after the addition of R6G^+^ to the cis compartment, a concentration-gradient forms along the z-axis above the bilayer with the following relative values:
- Successful reconstitution of Pdr5 into the planar bilayer was demonstrated by electrophysiological measurements (e.g. Figure 7). With the bilayer containing the reconstituted Pdr5 we were able to show that after the addition of R6G^+^ to the cis compartment in the presence of ATP/Mg^2+^cis, both the distribution of R6G^+^ between the compartments and the concentration of R6G^+^ in the individual compartments changed exponentially. This was only marginally due to due to the accumulation of the fluorescent dye in the membrane in control bilayer and Pdr5 bilayer. However, the determining factor for this kinetic course of the change in the distribution and concentration of R6G^+^ between the HLB compartments is the observation that Pdr5 obviously transports R6G^+^actively from the cis into the trans compartment.

| relative value | 1 | 4 | $\geq 10^3$ |
| --- | --- | --- | --- |
|  | *isotropic solution*<br>*($z \geq 10\mu\text{m}$)*<br>*high rotational*<br>*R6G<sup>+</sup>mobility*<br>*($r_0=0.02$)* | *bilayer surface*<br>*$z=+3\mu\text{m}$ to $+10\mu\text{m}$*<br>*partially restricted*<br>*R6G<sup>+</sup>mobility*<br>*($r_0=0.15$)* | *membrane plane*<br>*$z=0\mu\text{m}$ to $z=\pm 1\mu\text{m}$*<br>*largely immobilized R6G<sup>+</sup>*<br>*($r_0=0.28$)* |

## Conclusion

The above-described results show that the ABC transporter Pdr5 reconstituted in the HLB-system in the presence of ATP and Mg^2+^ cis, actively transports the fluorescent dye R6G^+^ from the cis compartment across the membrane, whereby the fluorescent dye first accumulates near the membrane surface and further accumulates in the membrane before the dye -is mediated by Pdr5 released from the membrane into solution on the trans side.

## CRediT authorship contribution statement

Daniel Blum: Data curation, Writing – original draft, Writing – review & editing. Lea-Marie Nentwig: Data curation, Writing – review & editing. Richard Wagner: Conceptualization, Formal analysis, Funding acquisition, Investigation, Methodology, Project administration, Resources, Supervision, Writing – review & editing. Lutz Schmitt: Conceptualization, Formal analysis, Funding acquisition, Investigation, Methodology, Project administration, Resources, Supervision, Writing – review & editing.

## DATA AVAILABILITY

Data will be made available on request.

## Acknowledgments

We thank Philip Bartsch (Constructor University, 2019) for the preparation of the cartoon in Figure 1B and Prof. Mathias Winterhalter (Constructor University) for support. DB was founded by the DFG Project Wa-681/15-1 and LMN was founded by the DFG Project Schm1279/17-1.

## Appendix Model: Horizontal Lipid Bilayer

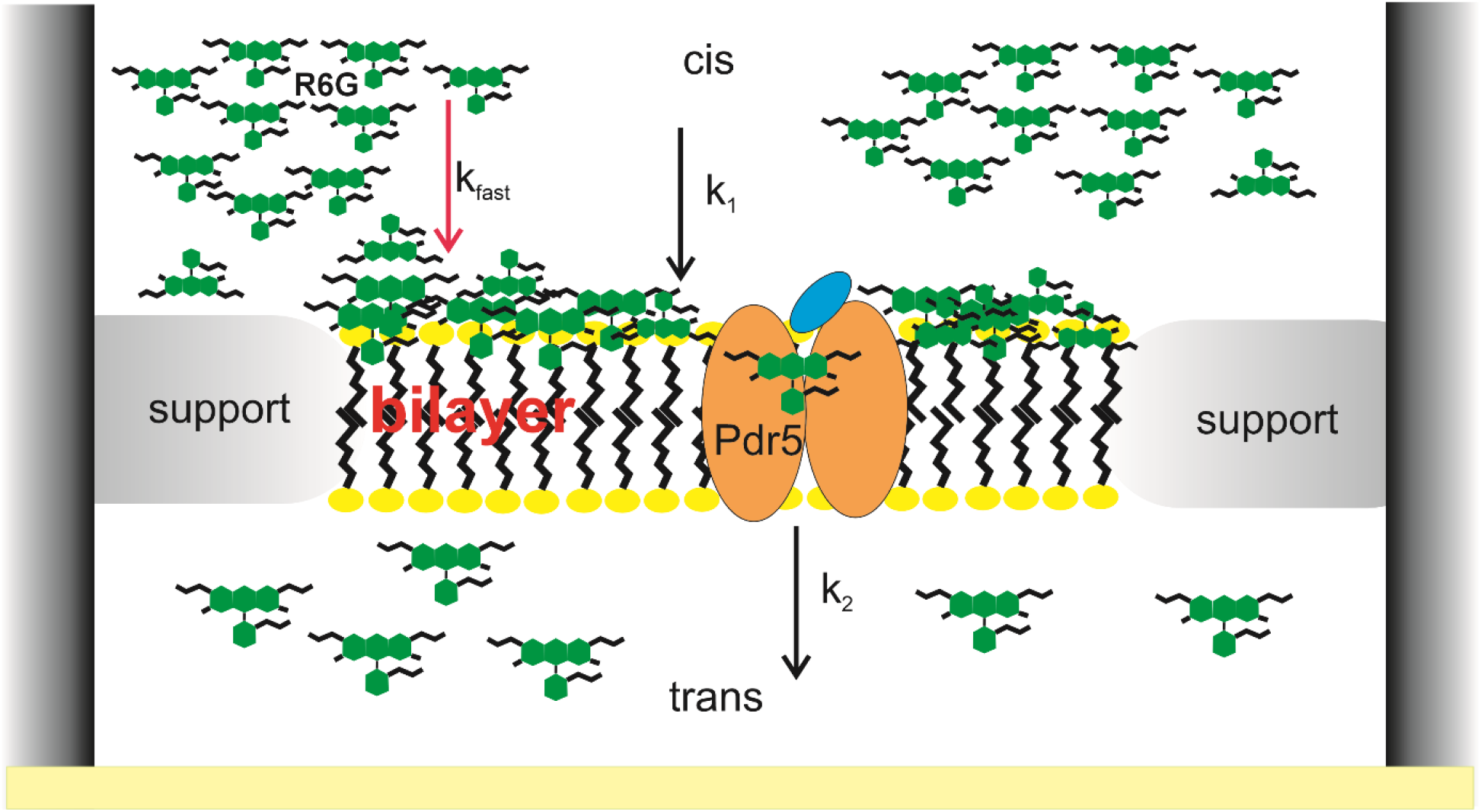

### Basic Observations

Rhodamine 6G when added to the cis compartment accumulates with regards to the local bilayer concentration very quickly by more than 3orders of magnitude on the membrane surface. Experimentally, this process could not be resolved in time (*k*_*fast*_ ≫ 7*min*^−1^). This effect is presumably due to the hydrophobic effect which, -according to the obtained data, revealed a R6G^+^ partition-coefficient of 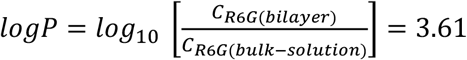. In the HLB-system with the bilayer containing the functional reconstituted ABC transporter Pdr5 and 5mM ATP/1mM Mg^2+^ (cis) this fast equilibration process was followed by a slower increase of the R6G^+^ concentration within the bilayer membrane plane with a rate constant of *k*_1_ = 0.74*min*^−1^. Concomitantly R6G^+^ is released in the trans compartment with a rate constant of *k*_2_= 0.43*min*^−1^. In a first approximation, the observed transport processes, which involved, -after the addition of 1μM R6G^+^ into the cis-compartment, the redistribution of R6G^+^ in the HLB -system, can be described by the following simplifying model:

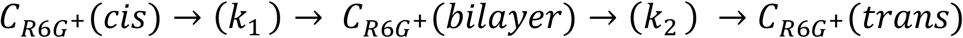

Now we can describe the time evolution of the R6G^+^ concentrations in different HLB compartments:

1. ***C***_***R*6*G***_^***+***^(***cis*** − ***surface***) **decays exponentially**

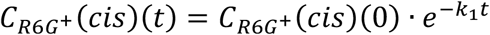
2. ***C***_***R*6*G***_^***+***^ (***bilayer***) **increases and decreases exponentially:**

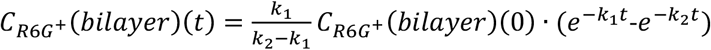
3. ***C***_***R*6*G***_^***+***^(***trans***) **increases exponentially:**

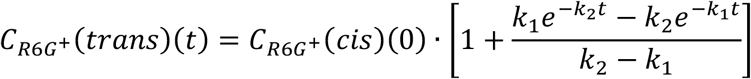

Now, neglecting the effect of the very fast accumulation of R6G^+^ at the membrane surface on the R6G^+^(cis) concentration, -which is valid due to the volume ratio of the cis compartment to the bilayer volume of about 10^6^, -we can plot the time course of the development of the R6G^+^ concentrations in the three HLB compartments after addition of 1μm R6G^+^ to the cis compartment. Where the bilayer contained the functional reconstituted ABC transporter Pdr5 and sufficient ATP/ Mg^2+^ (cis) to drive the Pdr5 mediated transport of R6G^+^ up to *t* → ∞.

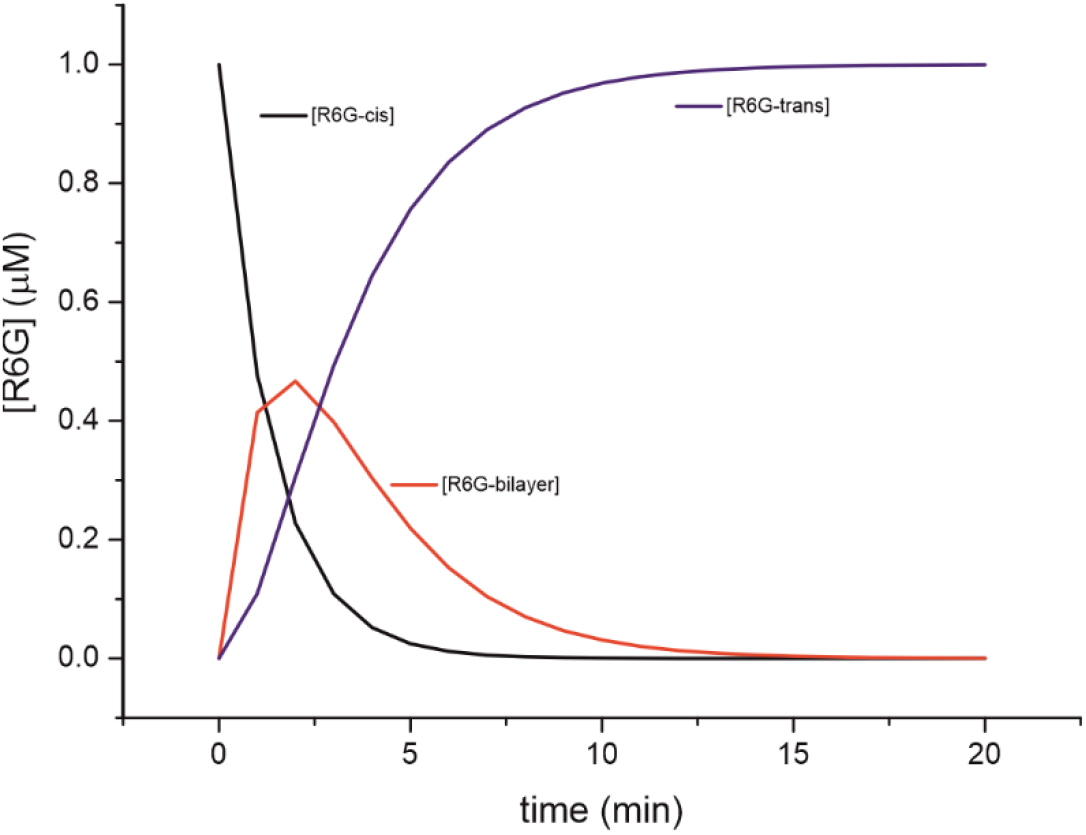

*Development of the R6G*^*+*^ *concentration in the 3 HLB compartments over time. The bilayer contained the reconstituted Pdr5 with 5mM ATP, 1mM Mg*^*2+*^ *(cis)*

It is obvious that the above simple, qualitative model does not provide any direct information about the molecular transport mechanisms of Pdr5, but the data and the model allow the kinetic classification of the Pdr5 transport capacity for active transport under the condition that sufficient energy (ATP/GTP) is available. Under this condition, the substrate R6G^+^ located on the cis-side would be completely transported to the trans compartment by Pdr5.

